# Closed-Loop Multi-Objective Optimization for Receptor-Selective Cell-Penetrating Peptide Design

**DOI:** 10.64898/2026.04.16.718169

**Authors:** Iori Yamahata, Teppei Shimamura, Shuto Hayashi

## Abstract

Cell-penetrating peptides (CPPs) can deliver diverse cargos into cells. However, designing CPPs with receptor-selective interaction profiles remains difficult because interactions with individual cell-surface components cannot be tuned independently. Here, we developed a closed-loop *in silico* framework for receptor-selective CPP design, in which receptor interactions are formulated as explicit objectives in a multi-objective optimization problem. We first constructed a CPP-like candidate library using a sequence generative model fine-tuned on known CPPs. The framework then evaluated candidate peptides by receptor-wise docking, molecular dynamics simulations, and MM/GBSA to compute receptor-wise binding scores. These scores were used iteratively to propose subsequent candidates by multi-objective Bayesian optimization. Applied to a CXCR4/NRP1 design setting, the framework identified candidates with more favorable predicted interaction profiles, characterized by higher CXCR4 binding scores and lower NRP1 binding scores. We selected 10 peptides from the computationally identified candidates for cell-based imaging and found that 4 showed higher enrichment in CXCR4-positive regions than in NRP1-positive regions under the tested conditions. These results show that the proposed framework provides a practical *in silico* approach for designing CPPs with receptor-selective interaction profiles.

## Introduction

Cell-penetrating peptides (CPPs) are promising molecules for intracellular delivery because they can transport diverse cargos, including proteins, nucleic acids, and small molecules, into cells [1, 2]. Their uptake involves multiple pathways and is often shaped by concurrent interactions with cell-surface receptors and glycans [3, 4]. Since interactions with individual cell-surface components cannot be tuned independently, it is difficult to achieve receptor-selective uptake and avoid off-target uptake [4, 5, 6].

Recent advances in CPP research include expanded sequence databases, uptake-related predictors, and deep-learning-based *in silico* screening and optimization methods [7, 8]. In parallel, selective CPP design has often relied on introducing or modifying sequence motifs to enhance interaction with a desired target molecule [9, 10, 11, 12, 13]. However, these approaches typically optimize CPP-likeness, uptake-related properties, or target interaction strength in isolation. Explicit treatment of target and non-target receptor interactions as separate design objectives remains limited. As a result, computational frameworks that evaluate receptor-wise interaction profiles and use them to guide sequence optimization under competing cell-surface interactions are still scarce.

Here, we present a closed-loop *in silico* framework for receptor-selective CPP discovery (Fig. 1), in which receptor interactions are formulated as explicit objectives in a multi-objective optimization problem. The resulting framework links CPP-like sequence generation, receptor-specific evaluation, and Bayesian optimization within a closed-loop design cycle, enabling candidate exploration based on receptor-interaction trade-offs rather than on a single-receptor objective.

**Figure 1.**
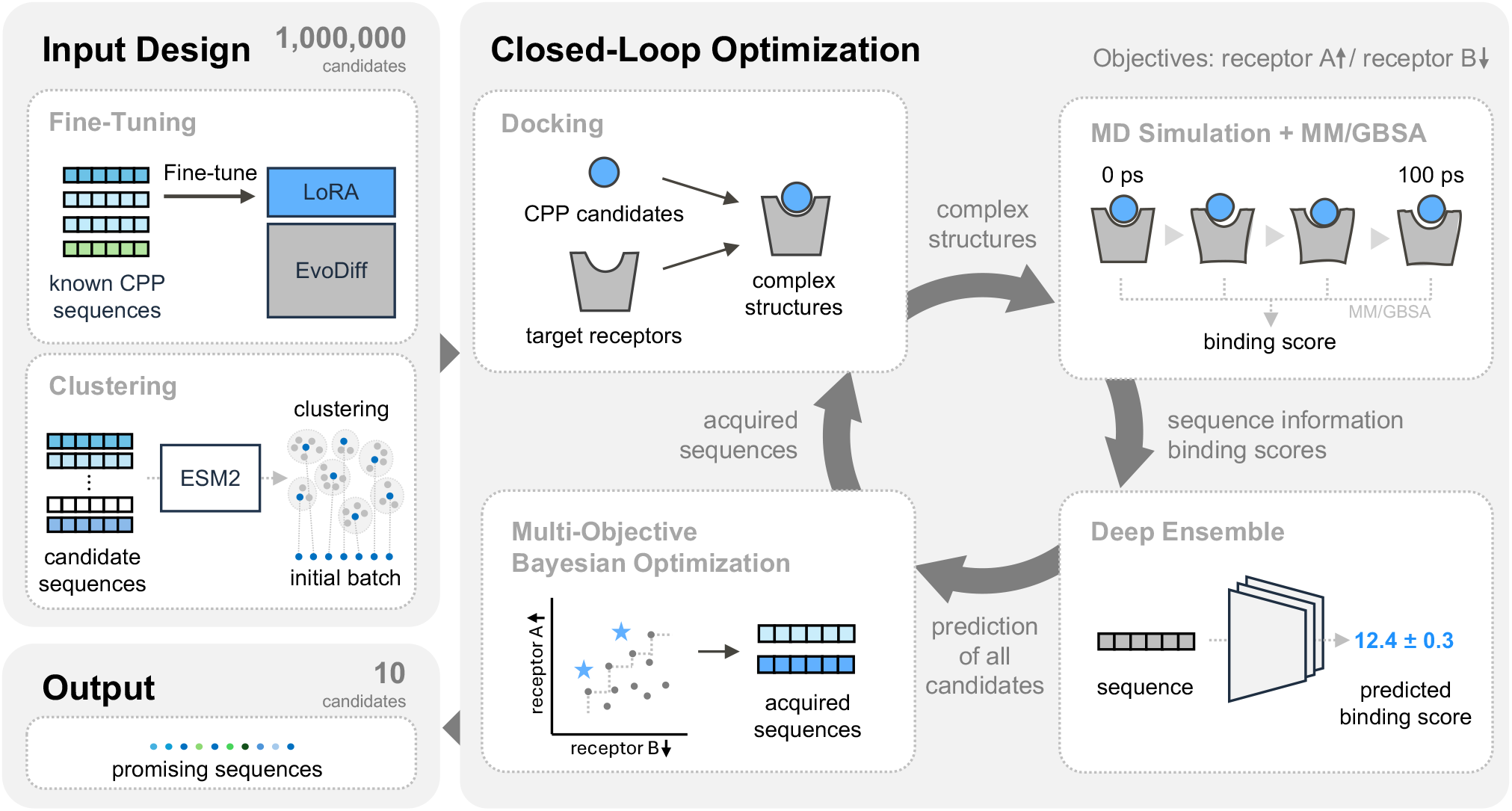
Schematic diagram of the closed-loop multi-objective optimization framework for receptor-selective CPP design. A total of 1,000,000 candidate sequences are generated using a sequence generative model (EvoDiff + LoRA) fine-tuned on known CPP sequences, and an initial candidate set is constructed by clustering feature vectors from a protein language model. In the optimization loop, receptor-wise binding scores are evaluated for selected peptide-receptor complexes using docking, MD simulations, and MM/GBSA. The deep ensemble surrogate model is then updated with the accumulated data, and multi-objective Bayesian optimization is used to propose sequences from the 1,000,000-sequence candidate library for evaluation in the next cycle. Through iterative repetition of this cycle, candidate peptides with receptor-selective predicted interaction profiles are identified.

The methodological contributions of this work can be summarized in three points. First, we formulate receptor-selective CPP design as a multi-objective optimization problem. By simultaneously treating the maximization of the target-receptor score and the minimization of the non-target-receptor score as independent objectives, we explicitly explore relative selectivity between receptors through candidates on the Pareto front. Such selectivity cannot be directly targeted in a single-objective formulation. Second, we construct a unified receptor-wise binding score. By combining docking-pose selection based on receptor-side interaction regions with MM/GBSA binding free energies normalized by the number of residues, we define an evaluation metric that is robust to variation in sequence length and binding location and that allows fair comparison across structurally distinct receptors. Third, we develop a closed-loop optimization architecture that makes computationally expensive physics-based evaluation practical at scale. A CPP-like sequence generative model based on a fine-tuned protein sequence generative model [14, 15], a deep ensemble surrogate model [16] that predicts the mean and variance of the receptor-wise binding score, and a noise- and batch-aware multi-objective Bayesian optimization acquisition function (qNEHVI) [17] are integrated into a single iterative cycle. Thus, the limited evaluation budget can be concentrated on the most promising candidates to be expected on the Pareto front.

We applied this framework to a CXCR4/NRP1 setting, in which CXCR4 was treated as the target receptor and NRP1 as the non-target receptor. This choice was motivated by the relevance of CXCR4 to leukemia-associated extramedullary infiltration and the well-established expression and functional roles of NRP1 in endothelial and other non-target cell contexts [4, 18, 19, 20]. In this setting, the framework identified candidates with more favorable predicted CXCR4/NRP1 interaction profiles. Confocal imaging of 10 computationally selected sequences in HeLa cells further revealed 4 of the 10 peptides with significantly higher enrichment in CXCR4-positive regions than in NRP1-positive regions. These results support closed-loop multi-objective *in silico* optimization as a useful framework for receptor-selective CPP design under competing cell-surface interaction constraints.

## Methods

### Overview

The overall framework is summarized in Fig. 1 and consists of an input-design phase followed by a closed-loop multi-objective optimization phase. In the input-design phase, a CPP-like candidate library is constructed by sampling from a sequence generative model that has been fine-tuned on known CPP sequences, and an initial subset of this library that broadly covers the candidate space is selected to seed the optimization phase. The closed-loop optimization phase then proceeds as a repeating cycle of four operations. Each candidate sequence is docked against both the target and the non-target receptors, and a receptor-wise binding score is computed by a short molecular dynamics simulation followed by MM/GBSA after pose filtering. The resulting sequence-score pairs are added to the training set of a deep ensemble surrogate model, which is retrained to predict the mean and variance of the binding score for each receptor. A multi-objective Bayesian optimization acquisition function then proposes the next batch of sequences from the candidate library for physics-based evaluation, and the proposed sequences enter the subsequent round of evaluation. The loop is terminated according to a stopping criterion based on the consistency of the top-ranked sequences, after which a final ranking is computed over all evaluated sequences, and a small set of top candidates is selected for downstream experimental validation.

The architectural choices outlined above are guided by three design principles. First, exploration is restricted to a CPP-like region of sequence space because the combinatorial space of amino-acid sequences is too large for random search or local motif grafting to cover the relevant region; we therefore use a generative model fine-tuned on known CPPs to encode CPP-likeness as an implicit prior over the candidate library. Second, the receptor-wise binding score used as an objective is constructed to be comparable across sequences of different lengths and across structurally distinct receptors. Docking poses are filtered against receptor-side interaction regions to ensure biological plausibility of the binding location. In addition, the MM/GBSA-derived binding free energy is normalized by the number of residues to remove length-dependent bias. Third, the cost of physics-based evaluation is allocated where it is most promising: a deep ensemble surrogate provides receptor-wise score predictions together with their uncertainties, and a noise- and batch-aware multi-objective acquisition function (qNEHVI) [17] uses these predictions to select sequences that are expected to expand the Pareto front in subsequent cycles. These three principles operate jointly within a single closed loop rather than in isolation, and each plays a distinct role in making receptor-selective CPP design tractable in practice. The remainder of this section describes each component of the framework in turn, beginning with the sequence generative model used to construct the CPP-like candidate library.

### CPP Sequence Generation Model and Candidate-Library Construction

#### Fine-tuning of the Sequence Generative Model

To enable efficient exploration in the vast peptide sequence space, a CPP-like candidate library is constructed by fine-tuning a sequence generative model on known CPPs. EvoDiff OADM640M [14], an autoregressive protein sequence generative model, is employed as the base model. A total of 1,082 CPP sequences from CellPPD [21, 22] are used as the training dataset. Fine-tuning is performed using Low-Rank Adaptation (LoRA) [15] on the CPP dataset. The fine-tuned model is then used to generate 100,000 sequences for each length from 9 to 18 residues, yielding a library of 1,000,000 CPP-like candidate sequences.

#### Selection of the Initial Batch for the Closed-Loop Optimization

An initial batch is selected from the library of the 1,000,000 candidate sequences to reduce redundancy while broadly covering the CPP-like candidate space. Embedding vectors are computed for each sequence using ESM2 (`esm2_t33_650M_UR50D`) [23], and *k*-means clustering is performed on residue-averaged embeddings. The number of clusters is set to 10,000, and the sequence with the smallest Euclidean distance to each cluster centroid is selected as its representative. The resulting 10,000 sequences are used as the initial set for the subsequent optimization loop.

### Closed-Loop Multi-Objective Optimization

The objective of the closed-loop optimization is to identify CPP candidates with favorable predicted score profiles from the pre-generated library of the 1,000,000 candidate sequences. In each cycle, selected candidates are evaluated, the prediction model is updated with the newly obtained data, and new candidate sequences are proposed for evaluation in the next cycle. As a practical stopping criterion, the loop is terminated when the set of the top 50 sequences remains unchanged for 10 consecutive cycles.

#### Docking and Selection of Docking Poses

To generate peptide-receptor complex structures, DiffDock [24] is used. Within each cycle, 20 docking poses are generated for each peptide-receptor pair. The poses are then filtered to remove physically or biologically unrealistic structures based on three criteria: intrapeptide clashes, peptide penetration into the receptor interior, and lack of contacts near the intended interaction region. Among the poses that pass all of the filters, the pose with the highest DiffDock confidence score is selected as the initial structure for subsequent MD simulations. The detailed filtering procedure and associated parameter settings are described in Supplementary Information.

#### MD Simulations and MM/GBSA

To evaluate each docked peptide-receptor complex in a physically relaxed state and assign a receptor-specific score for sequence selection, MD simulations followed by MM/GBSA analysis are performed. In the optimization loop, this score is used as the receptor-wise objective for each peptide-receptor pair.

For peptide-receptor complex structures obtained from docking, preprocessing is performed using the Protein Preparation Wizard in the Schrödinger suite [25]. Hydrogen bond networks are optimized, and protonation states of titratable residues are assigned at pH 7.0 using PROPKA [26, 27]. Terminal residues are capped as follows: ACE and NME groups are applied to the N- and C-termini of the receptor, respectively, while ACE and NH_2_ groups are applied to the CPP.

System topologies are generated using TLeap from AmberTools [28] with the AMBER ff14SB force field [29]. The complex is solvated using the TIP3P water model in a simulation box extending 12 Å beyond the outermost atoms of the complex. Na^+^ and Cl^−^ ions are added for charge neutralization, and the ionic strength is adjusted to 0.15 M. The resulting systems are converted to the GROMACS format using ACPYPE [30].

MD simulations are performed using GROMACS [31]. After energy minimization, the system is equilibrated for 100 ps under NVT while heating from 0 K to 310 K, followed by 100 ps equilibration under NPT at 310 K and 1 bar. A 100 ps production run is then performed under NPT at 310 K and 1 bar. A 2-fs timestep is used. Temperature and pressure are controlled using the V-rescale thermostat and the C-rescale barostat, respectively. Bonds involving hydrogen atoms are constrained using LINCS. Snapshots are saved every 10 ps during the production run.

To evaluate peptide-receptor interaction strength in the optimization loop, a binding score is defined from the MM/GBSA binding free energy computed using gmx_MMPBSA [32]. To reduce length-dependent bias across peptides of different lengths, the MM/GBSA value is normalized by the number of residues, and the sign is inverted so that larger values indicate stronger interaction:

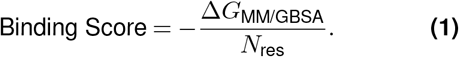

For MM/GBSA calculations, the internal dielectric constant is set to 2, the external dielectric constant to 80, and the salt concentration to 0.15 M.

#### Deep Ensemble Surrogate Model

To predict the receptor-wise binding scores and quantify their uncertainties for unevaluated sequences, a deep ensemble surrogate model is trained using the receptor-wise binding scores obtained as described above [16]. Each model takes a CPP sequence as input, maps it to a hidden representation through an embedding layer, and processes the representation using one-dimensional convolutional layers. The output layer predicts the mean and variance of the binding scores for each receptor. Three architectures with different convolutional configurations are used, and four models are trained for each architecture with different random initializations, yielding 12 models in total. Model training uses a negative log-likelihood loss, treating the MM/GBSA binding score as a sample from a normal distribution. Model specifications and training details are provided in Supplementary Information.

At each cycle, the deep ensemble is retrained using all binding score observations accumulated up to that point, and the mean and variance of the binding scores are predicted for the remaining unevaluated sequences.

#### Multi-objective Bayesian Optimization

To select sequences for MD simulations in the next cycle, multi-objective Bayesian optimization is applied using the deep ensemble surrogate model and qNEHVI (q Noisy Expected Hypervolume Improvement) [17]. qNEHVI seeks to expand the Pareto front by maximizing the expected hypervolume improvement while accounting for observation noise. This is appropriate in the present setting because the MM/GBSA-based scores vary owing to differences in initial structures and thermal fluctuations. In addition, qNEHVI allows for batch selection, which is suitable for large-scale parallel simulations. The reference point for qNEHVI is determined from the empirical distributions of the observed binding scores in the initial 10,000 evaluated sequences: *µ −* 2*σ* for the target receptor and *µ* + 2*σ* for the non-target receptor, where *µ* and *σ* denote the receptor-wise mean and standard deviation, respectively. In each cycle, qNEHVI is computed from the predictive distributions of the deep ensemble and used to select 2,000 unevaluated sequences for MD simulations in the next cycle. Details of the acquisition calculation are provided in Supplementary Information.

#### Selection of the Top Sequences

After the optimization loop, qNEHVI is computed using the trained deep ensemble while considering all simulated sequences as candidates. The top 10 sequences with the highest qNEHVI values are selected as candidates for experimental validation.

## Results

### Design of the CPP Exploration Space

To assess the effect of fine-tuning, we generated 1,000 sequences of length 9-18 residues from the default EvoDiff model and 1,000 from the fine-tuned model, and we computed the sequence embeddings using ESM2 (`esm2_t33_650M_UR50D`). We then defined a UMAP space using the embeddings from the default EvoDiff model and projected the default EvoDiff samples, fine-tuned samples, and known CPP sequences onto the same space (Fig. 2a). Samples from the default EvoDiff model were broadly distributed across the embedding space, whereas fine-tuned samples were concentrated closer to the region occupied by known CPPs.

**Figure 2.**
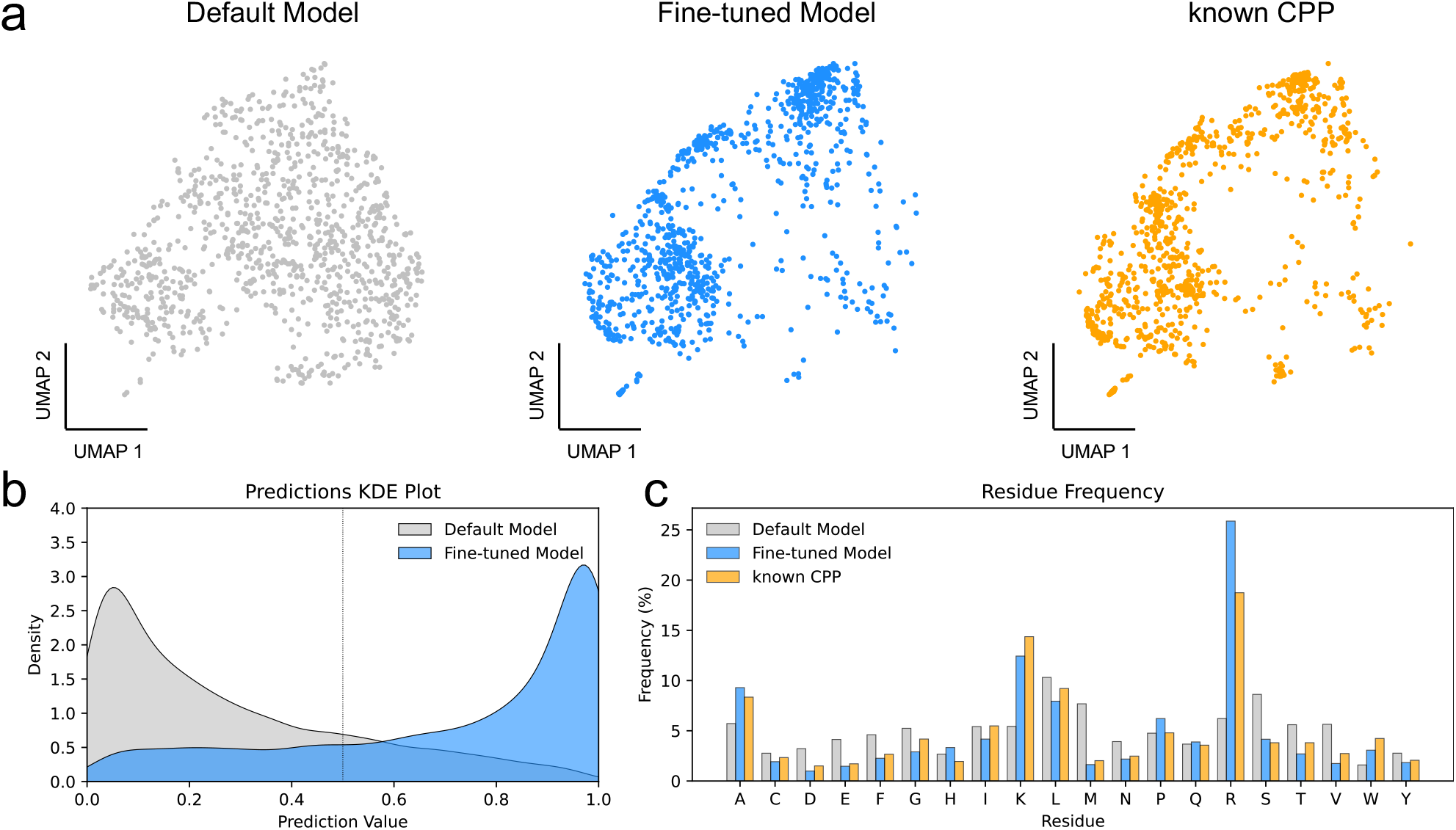
Construction of the CPP candidate library and sequence features before and after fine-tuning. (a) UMAP projection of ESM2 sequence embeddings. Left: default EvoDiff samples. Middle: fine-tuned samples. Right: known CPP sequences. (b) Kernel density estimation (KDE) of CPP-likeness scores estimated by the trained CPP/non-CPP classifier for default EvoDiff samples and fine-tuned samples. (c) Amino acid residue frequency distributions for default EvoDiff samples, fine-tuned samples, and known CPPs.

We next evaluated CPP-likeness of the generated sequences using two approaches: a trained CPP/non-CPP classifier and the existing CPP prediction tool c3pred [33]. The classifier was trained on data not used for fine-tuning; the data split, training procedure, and full performance metrics are described in Supplementary Information. The classifier achieved an AUC of 0.9148 on an independent test set. We then computed the CPP-likeness scores for the generated sequences using both the classifier and c3pred and compared the score distributions between default EvoDiff samples and fine-tuned samples (Fig. 2b; Fig. S3a,b). In both evaluations, fine-tuned samples shifted toward higher CPP-likeness than default EvoDiff samples.

We also compared the amino acid compositions between default EvoDiff samples and fine-tuned samples, using known CPPs as a reference. Relative to the default EvoDiff model, the fine-tuned model generated sequences enriched in basic residues such as arginine and lysine, while still retaining hydroxyl-containing and hydrophobic residues (Fig. 2c). Pairwise Tanimoto-distance distributions further indicated that the generated sequences retained sequence diversity (Fig. S3c). These results indicate that fine-tuning shifted the generated sequence distribution toward known CPP-like features while preserving sequence diversity. We therefore used the fine-tuned model to construct the candidate CPP library for subsequent receptor-wise evaluation and optimization.

### Initial Receptor-wise Binding Landscape

In this study, sequence design was formulated as a two-objective optimization problem in which CXCR4 was treated as the target receptor and NRP1 as the non-target receptor, with the aim of maximizing the CXCR4 binding score and minimizing the NRP1 binding score. We first defined the receptor-side interaction regions used for docking-pose filtering and subsequent receptor-wise scoring. We used PDB ID 3ODU for CXCR4 and PDB ID 7JJC for NRP1 (Fig. 3a).

**Figure 3.**
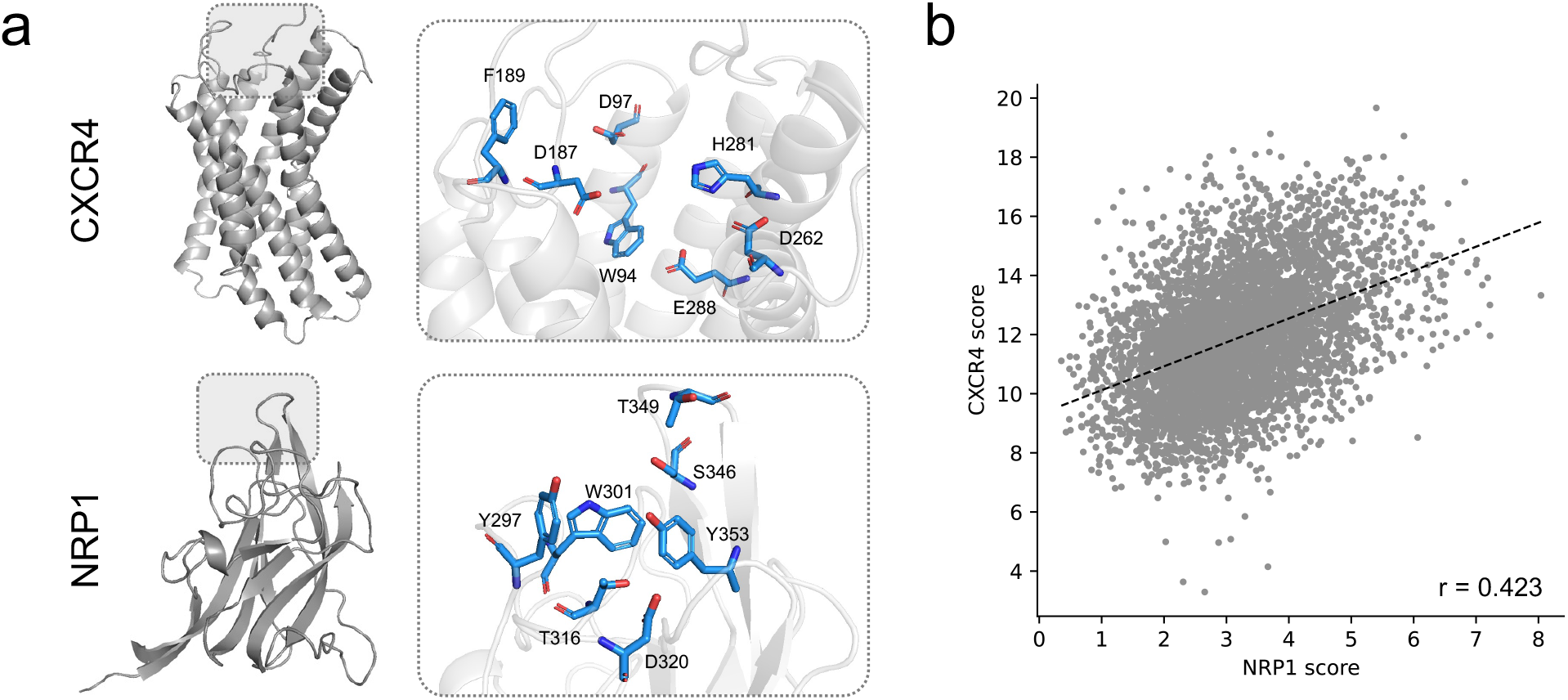
Docking target definition and correlation of receptor-wise binding scores. (a) Overall structures of the receptors. CXCR4 is shown on the left and NRP1 on the right. Receptor-side interaction regions used for docking-pose filtering are highlighted in blue. (b) Scatter plot of CXCR4 and NRP1 binding scores calculated for the initial candidate set of 10,000 sequences. The black dashed line indicates the linear regression line. Pearson’s correlation coefficient is *r* = 0.423 (*p* = 6.24 × 10^−226^).

Since each optimization cycle required thousands of evaluations, the optimization loop used a computationally tractable 100 ps MD protocol. Details of this protocol and its assessment against longer-timescale simulations are provided in Supplementary Information (Fig. S4).

Using this protocol, we docked the initial 10,000 candidate sequences against both receptors and computed receptor-wise binding scores by MD simulations and MM/GBSA for complexes that passed the pose filtering. Across the initial set, CXCR4 and NRP1 binding scores were positively correlated (Fig. 3b), indicating that improvement in the CXCR4 binding score could unintendedly enhance the NRP1 binding score.

### Exploration of the Receptor-Selective CPP Candidates by Multi-Objective Bayesian Optimization

We performed multi-objective Bayesian optimization over the candidate CPP library using receptor-wise binding scores as objectives.

To monitor surrogate-model performance during optimization, we calculated the correlation between the predicted binding scores and the observed MM/GBSA binding scores on a validation subset. In each optimization cycle, the available evaluated sequences were randomly split into training and validation subsets at a ratio of 9:1, and the surrogate models were retrained on the training subset.

This correlation increased over the course of optimization (Fig. 4a), indicating progressive improvement in the agreement between predicted and observed scores.

**Figure 4.**
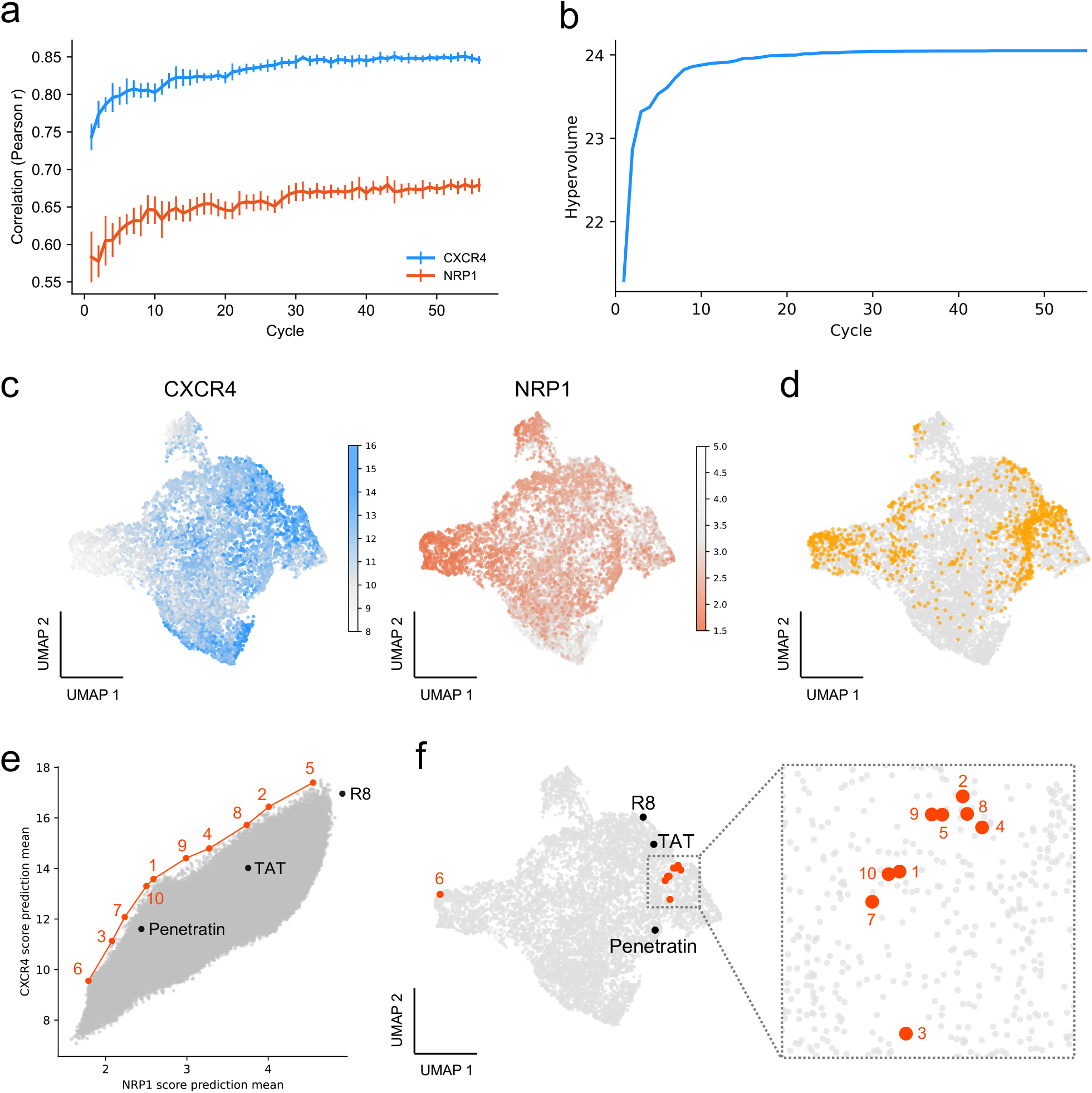
Exploration process and sequence distribution in multi-objective optimization. (a) Correlation coefficient between the deep ensemble mean predictions and the MM/GBSA binding scores on a validation subset in each optimization cycle. (b) Hypervolume values computed from predicted mean binding scores in each cycle using the deep ensemble model trained in the final cycle. (c) UMAP projection of ESM2 sequence embeddings colored by predicted binding scores. Left: CXCR4 predicted binding scores. Right: NRP1 predicted binding scores. (d) UMAP projection of the same embeddings highlighting sequences proposed by Bayesian optimization (yellow points). (e) Scatter plot of CXCR4 and NRP1 binding scores predicted by the surrogate model (gray points), with representative known CPPs and the selected sequences overlaid. The selected sequences are shown in red and numbered 1-10 (corresponding to S-01–S-10). (f) UMAP projection of all sequences, with the selected sequences shown in red and representative known CPPs shown in black.

To quantify the optimization progress, we performed a post-hoc hypervolume analysis using predicted mean binding scores. Sequences obtained in each cycle were re-evaluated using the deep ensemble model trained in the final cycle, and the hypervolume was calculated relative to the reference point. Hypervolume increased in early cycles and then approached a plateau in later cycles (Fig. 4b).

To examine which regions of the sequence space were preferentially explored, we visualized surrogate-predicted binding scores on the ESM2/UMAP embedding space in the final cycle (Fig. 4c) and overlaid the sequences proposed by Bayesian optimization on the same space (Fig. 4d). The proposed sequences were concentrated in regions associated with higher predicted CXCR4 scores and lower predicted NRP1 scores.

After completing the optimization loop, we selected 10 top-ranked sequences, denoted S-01 to S-10, from the set of candidates evaluated by MD simulations. Table 1 summarizes their sequences, lengths, and predicted receptor-wise binding scores together with those of three representative known CPPs used for comparison in subsequent analysis: TAT, R8, and Penetratin.

**Table 1.**
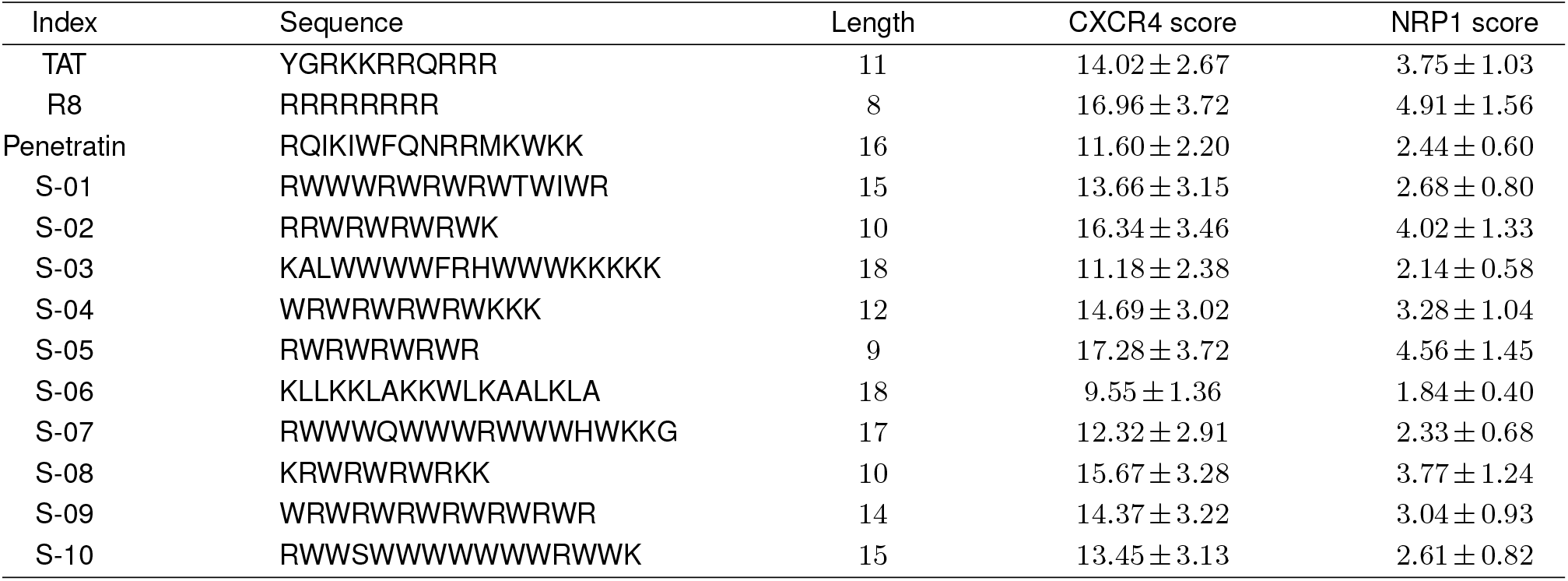
Amino acid sequences, lengths, and predicted binding scores for representative known CPPs (TAT, R8, Penetratin) and selected sequences (S-01 to S-10).

We then plotted the predicted CXCR4 and NRP1 scores for all candidate sequences and overlaid S-01 to S-10 together with representative known CPPs (Fig. 4e). These sequences showed favorable predicted CXCR4/NRP1 score combinations, consistent with the optimization objectives defined in this study. We also projected them onto the UMAP embedding relative to the full candidate library and to known CPPs (Fig. 4f). We found that one of them, S-09 (WRWRWRWRWRWRWR), was present in the fine-tuning dataset.

### Cell-Based Evaluation of Receptor-Associated Peptide Enrichment

To examine whether the selected sequences showed receptor-associated localization patterns consistent with the *in silico* predictions, we performed cell-based evaluation using confocal microscopy with fluorescently labeled peptides (TAT, R8, Penetratin, and selected peptides) and quantified the peptide enrichment relative to receptor-positive regions. HeLa cells were used as a tractable model system because both CXCR4 and NRP1 are expressed in HeLa cells and can be evaluated in the same cellular background under a common imaging workflow. Cells were incubated individually for 1 hour with each fluorescently labeled peptide, followed by immunofluorescence staining for CXCR4 or NRP1.

To quantify receptor-associated localization, we analyzed the images using the procedure described in Supplementary Information. Briefly, peptide-positive ROIs were defined from the peptide channel, receptor-positive ROIs were defined from the receptor channel, and an enrichment score was calculated as the ratio of mean peptide intensity in receptor-positive ROIs to that in peptide-positive ROIs.

We first quantified and summarized the mean enrichment scores of each peptide with respect to CXCR4-positive and NRP1-positive regions (Fig. 5a). TAT, R8, S-07, S-09, and S-10 were excluded from this comparison because fluorescence intensity was insufficient for reliable quantification under the imaging conditions used. Among the quantified peptides, S-01, S-02, S-03, and S-08 showed higher enrichment scores with respect to CXCR4-positive regions than NRP1-positive regions. Also, Penetratin showed the same tendency.

**Figure 5.**
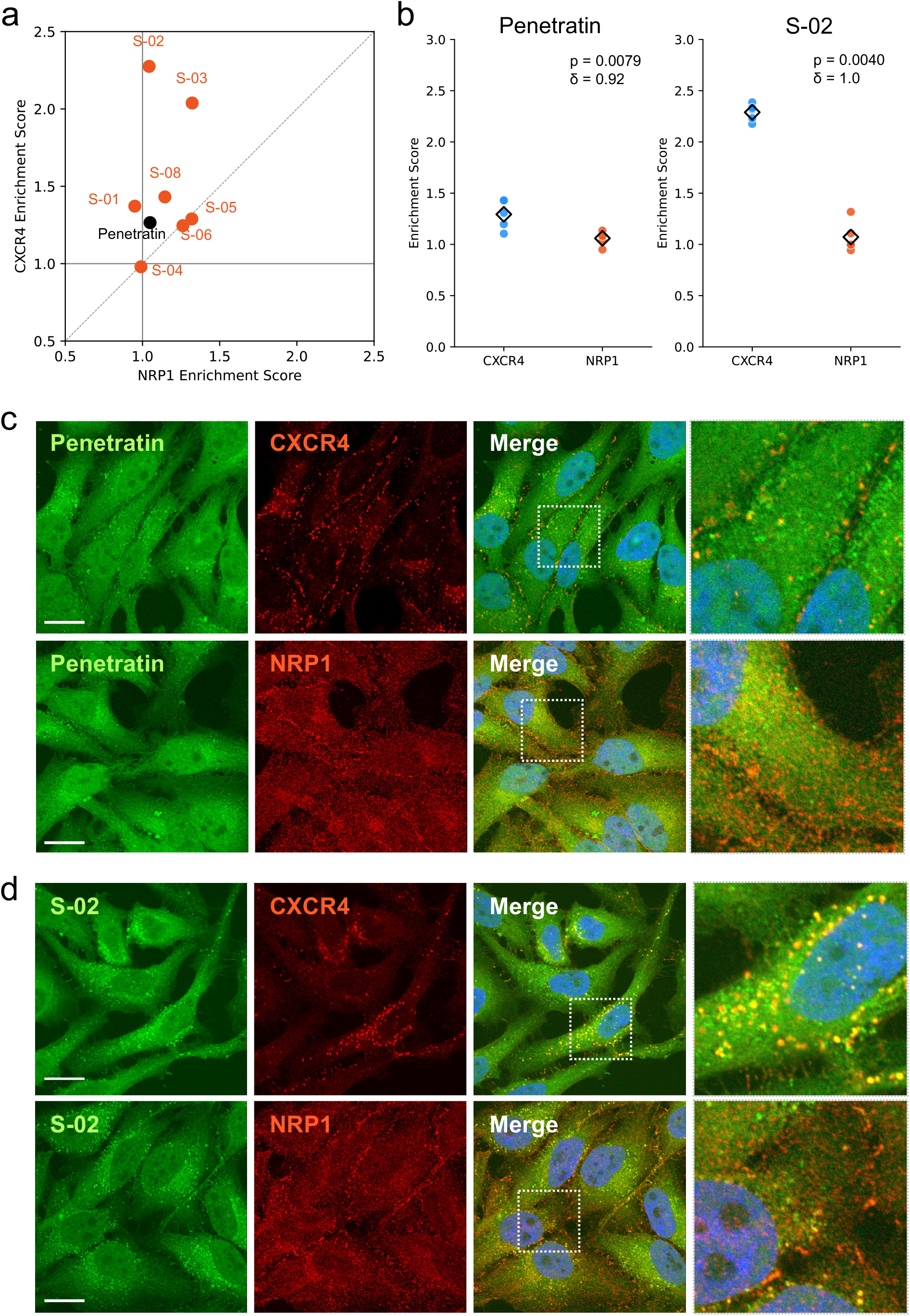
Cell-based evaluation of receptor-associated peptide enrichment. (a) Mean enrichment scores for quantified peptides with respect to CXCR4-positive and NRP1-positive regions. TAT, R8, S-07, S-09, and S-10 were excluded because fluorescence intensity was insufficient for reliable quantification. (b) Enrichment score distributions for Penetratin and S-02. *P* values were computed using a one-sided Mann-Whitney *U* test, and the corresponding effect sizes are indicated as Cliff’s *δ*. (c, d) Representative confocal images of Penetratin (c) and S-02 (d) under CXCR4 or NRP1 immunostaining. Green: peptide; red: receptor; blue: DAPI. Scale bar: 20 µm.

As representative examples from this analysis, the enrichment score distributions for Penetratin and S-02 are shown in Fig. 5b. For both Penetratin and S-02, enrichment scores were significantly higher with respect to CXCR4-positive regions than NRP1-positive regions according to a one-sided Mann-Whitney *U* test, and the corresponding effect sizes are indicated as Cliff’s *δ* (Fig. 5b). Representative confocal images showed punctate intracellular signals for both peptides, with Penetratin exhibiting partial overlap with CXCR4-positive regions and S-02 showing a qualitatively stronger tendency toward CXCR4-positive regions (Fig. 5c,d). The enrichment score distributions for the other quantified peptides are provided in Fig. S7. These results indicate that several selected peptides showed higher enrichment in CXCR4-positive regions than NRP1-positive regions under the tested conditions.

## Discussion

In this study, we developed a closed-loop multi-objective optimization framework for designing CPPs with receptor-selective interaction profiles. Specifically, we formulated the design problem as a multi-objective optimization over receptor-wise binding scores and solved it through a closed-loop *in silico* workflow. In the CXCR4/NRP1 case study, CXCR4 and NRP1 binding scores were positively correlated across the initial candidate set, indicating that sequences with stronger interaction with CXCR4 also tended to show stronger interaction with NRP1. This observation directly motivates the multi-objective formulation: a single-objective strategy that maximizes CXCR4 interaction alone would also tend to enrich peptides that interact strongly with NRP1, making receptor-selective design difficult to achieve. Despite this inherent trade-off, the optimization process identified candidate peptides with improved predicted CXCR4-to-NRP1 score balance. Cell-based imaging further provided supporting evidence that several of these candidates were enriched in CXCR4-positive regions to a greater extent than in NRP1-positive regions. These findings support the value of formulating receptor-selective CPP design under competing receptor-interaction constraints as an explicit multi-objective optimization problem.

Three methodological aspects of this framework are particularly important in this context. First, the multi-objective formulation itself makes it possible to express receptor selectivity as an explicit design objective rather than as a property to be hoped for downstream; the output of optimization is therefore expected to be a set of candidates lying on the trade-off front between target and non-target receptor interactions, from which sequences biased toward the target side can be selected directly. Second, the receptor-wise binding score used as the optimization objective combines receptor-side pose filtering with length-normalized MM/GBSA, which allows peptides of different lengths and structurally distinct receptors to be compared on a common scale and reduces the risk that the optimizer would simply favor longer sequences or biologically implausible binding poses. Third, the closed-loop optimization architecture is particularly important for making broader exploration of CPP-like sequence space more practical. This architecture combines a CPP-like generative model [14, 15], a deep ensemble surrogate [16], and a noise-aware multi-objective acquisition function (qNEHVI) [17]. Rather than restricting design to the local sequence changes typical of grafting-style or motif-based strategies [9, 11, 12, 13], the framework concentrates computationally expensive receptor-wise evaluation on candidates most likely to improve the multi-objective trade-off, without requiring exhaustive screening.

The potential utility of this framework is not limited to the CXCR4/NRP1 case examined here. In principle, other receptor-selective design settings can also be addressed within the same framework. This is because neither the receptor-wise scoring procedure nor the closed-loop optimization is specific to a particular receptor structure; only the receptor pair and the associated optimization objectives need to be redefined. Moreover, the number of objectives is not restricted to two. Cytotoxicity and aggregation propensity, for example, are both relevant constraints in the development of peptide-based drug delivery systems (DDSs). CPPs can show sequence- and cargo-dependent cytotoxicity through strengthened nonspecific interactions with cellular membranes [34, 35], and aggregation is a well-recognized developability challenge for peptide and protein therapeutics [36]. In principle, cytotoxicity could be incorporated using peptide-toxicity predictors or task-specific models trained on annotated datasets, whereas aggregation propensity could be incorporated using sequence-based aggregation predictors. Extending the framework in this way would add further objectives to the existing multi-objective formulation without requiring changes to the underlying optimization architecture, and could enable the search for candidates that balance receptor-associated interaction profiles with practical developability requirements.

A central limitation of the present study is that the framework was optimized against computational receptor-wise scores rather than against biological selectivity itself. The current framework operates on a computational proxy: the multi-objective formulation treats receptor-wise scores as the design objective, the binding score itself is a physics-based approximation derived from a short molecular dynamics simulation followed by MM/GBSA, and the closed-loop optimization explores the trade-off front defined by these computed scores rather than the trade-off front defined by measured cellular phenotypes. Receptor-associated enrichment in cell-based imaging provides supporting evidence that the identified peptides captured biologically relevant differences, but this readout should not be regarded as a complete measure of selectivity. In practice, biological selectivity is also influenced by processes beyond receptor interaction strength and local enrichment, including membrane perturbation, endocytic routing, intracellular trafficking, and endosomal escape [2, 5, 37, 38]. A natural next step is therefore to incorporate experimental readouts such as cellular uptake, intracellular localization, or receptor-associated enrichment directly into the design loop, combined with the computational proxy. This would consider not only optimizing computational receptor-wise proxies but also optimizing experimentally observed cellular phenotypes. The present study establishes a starting point for moving receptor-selective CPP design from *in silico* optimization based on proxy objectives toward closed-loop optimization of experimentally defined cellular function.

## Data Availability

The fine-tuning code and trained weights for the sequence generative model, the optimization-cycle pipeline code, and the Fiji macro used for image quantification are available on GitHub (https://github.com/IoriYamahata/CPP_Design). Note that the provided source code is a simplified version intended for concept verification; specific implementations for large-scale parallel computing (multi-node execution) used in the supercomputing environment have been omitted to ensure portability. All other data supporting the findings of this study are included in the main text and the Supplementary Information.

## ACKNOWLEDGEMENTS

This work was supported by JSPS KAKENHI Grant Numbers JP22K17993, JP24H01756, and JP25H01213, JST PRESTO Grant Number JPMJPR24TA, JST BOOST Program Grant Number JPMJBY24G3, the Takeda Science Foundation, and Project Mirai Cancer Research Grants (to S.H.); and JSPS KAKENHI Grant Numbers JP22H04925, JP23H04938, and JP26K03026, AMED Grant Numbers JP25gm2010002, JP25nk0101112, JP25wm0625519, JP25wm0325068, JP25zf0127012, and JP25tm0424226, JST Moonshot R&D Grant Number JPMJMS2025, the Medical Research Center Initiative for High Depth Omics (Institute of Science Tokyo), Multilayered Stress Diseases (Institute of Science Tokyo), and the Uehara Memorial Foundation (to T.S.). This work was also supported in part by JST SPRING, Grant Number JPMJSP2125 (Nagoya University Make New Standard Next-Generation Researcher Program), and the Nagoya University CIBoG WISE Program (to I.Y.). This study was carried out using the TSUBAME supercomputer at Institute of Science Tokyo.

## AUTHOR CONTRIBUTIONS

IY: Conceptualization, Investigation, Formal analysis, Software, Visualization, Writing - original draft.

TS: Methodology, Supervision, Writing - review & editing, Funding acquisition. SH: Methodology, Supervision, Writing - review & editing, Funding acquisition.

## COMPETING FINANCIAL INTERESTS

The authors declare no conflict of interest.

## Supplementary Information

### Closed-Loop Multi-Objective Optimization for Receptor-Selective Cell-Penetrating Peptide Design

Iori Yamahata, Teppei Shimamura, Shuto Hayashi

#### Fine-tuning of the Sequence Generative Model

To adapt EvoDiff OADM640M to the CPP sequence distribution, EvoDiff is fine-tuned on experimentally reported CPP sequences using Low-Rank Adaptation (LoRA), in which trainable low-rank matrices are introduced while the pretrained model weights are kept fixed.

In this study, LoRA is applied to the MaskedConv1d layers in EvoDiff, which capture local sequence features through one-dimensional convolutions. The convolution kernel size is 5, and the LoRA hyperparameters are set to *r* = 8, *α* = 2, and dropout = 0.0.

Fine-tuning is performed on the 1,082 CPP sequences collected from CellPPD, as described in the main text. All sequences are converted to uppercase amino-acid strings, and the dataset is randomly split into training and validation sets in a 9:1 ratio. Training uses the OAMaskedCrossEntropyLoss implemented in EvoDiff and the Adam optimizer with a weight decay of 0. An exponential learning-rate schedule is applied using a LambdaLR scheduler: the learning rate is increased exponentially from 1 × 10^−7^ to 1 × 10^−4^ during the first 10% of epochs, held at 1×10^−4^ until 50% of epochs, and then decayed exponentially to 1 × 10^−7^ over the remaining epochs. Fine-tuning is run for 3,000 epochs with a batch size of 100, and the final adapted model is used for sequence generation.

#### Training of the CPP Classifier

To assess whether the fine-tuned generative model produces sequences with CPP-like characteristics, we train a binary classifier to distinguish CPPs from non-CPPs using the dataset reported by Park et al. [1]. The development dataset consists of 561 CPP sequences and 1,097 non-CPP sequences, none of which are used for fine-tuning the generative model. These sequences are split into training and validation sets in an 8:2 ratio by stratified random sampling based on the CPP/non-CPP labels, using a fixed random seed. For final evaluation, we use the independent test set provided by Park et al., which consists of 150 CPP and 150 non-CPP sequences.

Input sequences are integer-encoded using a 20-amino-acid vocabulary with an additional PAD token. Sequences are right-padded, and the maximum sequence length is set to the maximum length observed in the development dataset. The classifier consists of an embedding layer, two one-dimensional convolutional layers with kernel sizes of 3 and 5, each followed by batch normalization and dropout, and a fully connected output layer. The embedding dimension and hidden dimension are both set to 16, and the dropout rate is 0.2.

The model is trained for 1,000 epochs with a batch size of 16 using binary cross-entropy loss with logits and the Adam optimizer with a learning rate of 1 × 10^−5^. The model checkpoint with the lowest validation loss is selected for final evaluation. On the independent test set, each sequence is classified as CPP when the predicted probability exceeds 0.5 and as non-CPP otherwise. The classifier achieved an AUC of 0.9148, an accuracy of 84.00%, a precision of 0.8643, a recall of 0.8067, and an F1 score of 0.8345. The trained classifier is then applied to the generated sequences to estimate the probability that each sequence is CPP-like. The architecture of the classifier is shown in Fig. S1.

#### Selection of Docking Poses

To define an initial structure for MD simulation from the multiple poses generated by DiffDock, 20 docking poses are prepared for each peptide-receptor pair and filtered using the criteria below.

1. **Exclusion of poses with intrapeptide clashes**. If at least one pair of non-covalently bonded atoms in the peptide has an interatomic distance of 1.5 Å or less, the pose is considered to contain an intrapeptide clash and is excluded.
2. **Exclusion of poses with peptide penetration into the receptor interior**. To detect poses in which the peptide unnaturally penetrates into the receptor interior, we use the *S* function from DiffBP:

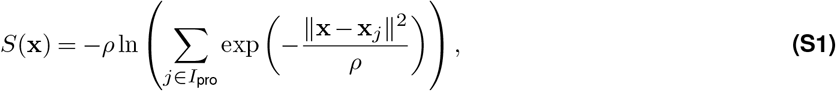

where **x** denotes the 3D coordinates of a peptide atom, **x**_*j*_ denotes the 3D coordinates of the *j*-th receptor atom, and *I*_pro_ denotes the index set of all receptor atoms. A pose is excluded if at least one peptide atom satisfies

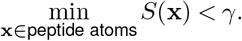

The hyperparameters are set to *ρ* = 0.3 and *γ* = 0.1, respectively.
3. **Exclusion of poses lacking contacts near the intended interaction region**. To exclude poses located far from the receptor-side interaction region of interest, receptor-specific key residues are defined, and at least one peptide atom is required to contact this region. A pose is excluded if no peptide atom is within 4.2 Å of any atom in the corresponding receptor-specific key residues. For CXCR4, the key residues are W94, D97, D187, F189, D262, H281, and E288 based on prior structural and functional analyses of the receptor [2]. For NRP1, the key residues are Y297, W301, T316, D320, S346, T349, and Y353 based on structural analysis of the CendR binding pocket [3]. Among the poses that pass all three filters, the one with the highest DiffDock confidence score is selected as the initial structure for subsequent MD simulations. If no pose passes all filters for a given peptide-receptor pair, that pair is excluded from subsequent MD simulations.

#### Validation of the 100 ps MD Protocol

Since each optimization cycle requires thousands of evaluations, the main optimization loop employed a computationally tractable 100 ps MD protocol. To assess whether this short-timescale protocol was adequate for sequence selection, we compared the 100 ps condition with a longer 100 ns reference condition using the 10 peptides selected from candidates evaluated during the optimization loop (S-01 to S-10), each evaluated against both CXCR4 and NRP1. For each peptide-receptor pair, 10 simulations were initiated from 10 different initial structures generated by DiffDock. The 100 ps simulations were performed using the same protocol as that adopted in the main optimization loop.

For the 100 ns reference condition, CXCR4, a transmembrane receptor, was simulated in a membrane-embedded system, whereas NRP1, treated as an ectodomain receptor, was simulated in aqueous solution without an explicit membrane. For CXCR4, the membrane orientation and position relative to the protein were estimated using the OPM/PPM server [4, 5], and a bilayer composed of 30% cholesterol and 70% POPC was constructed using the Membrane Builder in CHARMM-GUI [6, 7]. The force field, water model, ion concentration, and other simulation settings were the same as those used in the main MD protocol. Binding scores for the 100 ns simulations were calculated by MM/GBSA analysis using 100 frames sampled at intervals of 1,000 frames from each trajectory. Simulations with sign-inverted MM/GBSA-derived energy values greater than 100 before normalization by the number of residues were regarded as crashed trajectories and were excluded from subsequent analysis. For both timescales, MM/GBSA scores were averaged across the initial structures to obtain one score for each peptide-receptor pair, and Pearson and Spearman correlations between the 100 ps and 100 ns scores were calculated for each receptor. The resulting 100 ps and 100 ns scores showed a positive correlation for both receptors (Fig. S4).

#### Deep Ensemble Model Architecture and Training

To predict receptor-wise binding scores from peptide sequences during the optimization loop, deep ensemble surrogate models are trained. The models took peptide sequences as input and output the mean and variance of the predicted binding score for each receptor. The architecture of the surrogate model is illustrated in Fig. S2. Input sequences were encoded as integer amino-acid indices using the amino-acid dictionary. They were then right-padded with the PAD token to the maximum sequence length defined in the model configuration, and sequences longer than this length were truncated. The encoded sequence was mapped to a hidden representation through an embedding layer and then processed by repeated one-dimensional convolutional blocks to extract local sequence features. These features were flattened into a fixed-length representation and passed to regression heads that output the mean and variance of the binding score for each receptor, with the variance output constrained to be positive using the softplus function.

Using this basic structure, three model configurations with different convolutional settings were prepared. For each configuration, four models were trained with different random initializations, yielding 12 models in total. Architecture-specific hyperparameters are summarized in Table S1.

Before training, sequences were excluded if any receptor-wise sign-inverted MM/GBSA-derived energy value exceeded a predefined threshold before normalization by the number of residues. Such cases were regarded as crashed trajectories. In each optimization cycle, the available dataset was randomly split into training and validation subsets at a 9:1 ratio. All models were trained with a batch size of 500 and a maximum sequence length of 18 using Gaussian negative log-likelihood loss and the Adam optimizer with a weight decay of 1× 10^−5^. Training was performed for up to 5,000 epochs. The checkpoint with the lowest validation loss was retained, and early stopping was applied based on validation loss using the patience value specified for each model configuration. The predictive mean and variance used in downstream acquisition were obtained by aggregating the outputs of the 12 ensemble members.

#### Deep Ensemble Model Architecture and Training

To predict receptor-wise binding scores from peptide sequences during the optimization loop, deep ensemble surrogate models are trained. The models take peptide sequences as input and output the mean and variance of the predicted binding score for each receptor. The architecture of the surrogate model is illustrated in Fig. S2. Input sequences are encoded as integer amino-acid indices using the amino-acid dictionary. They are then right-padded with the PAD token to the maximum sequence length defined in the model configuration, and sequences longer than this length are truncated. The encoded sequence is mapped to a hidden representation through an embedding layer and then processed by repeated one-dimensional convolutional blocks to extract local sequence features. These features are flattened into a fixed-length representation and passed to regression heads that output the mean and variance of the binding score for each receptor, with the variance output constrained to be positive using the softplus function.

Using this basic structure, three model configurations with different convolutional settings are employed. For each configuration, four models are trained with different random initializations, yielding 12 models in total. Architecture-specific hyperparameters are summarized in Table S1.

Before training, sequences are excluded if any receptor-wise sign-inverted MM/GBSA-derived energy value exceeds a predefined threshold before normalization by the number of residues. Such cases are regarded as crashed trajectories. In each optimization cycle, the available dataset is randomly split into training and validation subsets at a 9:1 ratio. All models are trained with a batch size of 500 and a maximum sequence length of 18 using Gaussian negative log-likelihood loss and the Adam optimizer with a weight decay of 1× 10^−5^. Training is performed for up to 5,000 epochs. The checkpoint with the lowest validation loss is retained, and early stopping is applied based on validation loss using the patience value specified for each model configuration. The predictive mean and variance used in downstream acquisition are obtained by aggregating the outputs of the 12 ensemble members.

#### Details of Multi-objective Bayesian Optimization

To select peptide sequences for MD simulation in each optimization cycle, we use a deep-ensemble-based qNEHVI acquisition function [8]. Since qNEHVI is formulated for maximization, the two-objective vector for each sequence *x* is defined as

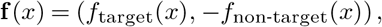

where *f*_target_(*x*) and *f*_non-target_(*x*) denote the actual binding scores for the target and non-target receptors, respectively.

Let *M* = 12 denote the number of surrogate models in the deep ensemble. For each surrogate model *m*, receptor-wise predictive means are computed for both previously evaluated sequences and unevaluated candidate sequences. The model-wise predictive mean vector for sequence *x* is defined as

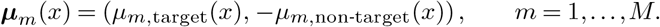

In the acquisition calculation, the model-wise predictive mean vectors are treated as ensemble-derived samples of the objective landscape. Accordingly, for each surrogate model *m*, a model-specific Pareto set is constructed from the predicted objective vectors of the previously evaluated sequences,

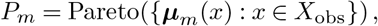

where *X*_obs_ denotes the set of previously evaluated sequences. The variance outputs of the surrogate models are not used in the acquisition calculation.

For a candidate batch *X*_cand_, the model-specific hypervolume improvement is defined as

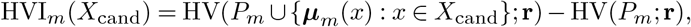

where **r** denotes the reference point and HV(*·*; **r**) denotes the hypervolume measured with respect to **r**. The acquisition value is defined as the expectation of the model-specific hypervolume improvement under the empirical distribution over ensemble members:

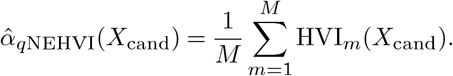

The reference point **r** is defined from the empirical distributions of the observed binding scores in the initial 10,000 evaluated sequences. Specifically, the reference point is set to *µ −* 2*σ* for the target receptor and *µ* + 2*σ* for the non-target receptor, where *µ* and *σ* denote the receptor-specific mean and standard deviation, respectively. In each cycle, acquisition values are computed for previously unevaluated sequences, and the top 2,000 sequences are selected for MD simulation in the next cycle.

#### Peptide Synthesis

Candidate peptide sequences were chemically synthesized by Scrum Inc. (Tokyo, Japan). To enable fluorescence visualization, the N-terminus of each peptide was labeled with 6-carboxyfluorescein (6-FAM), and the C-terminus was amidated. The purity of all synthesized peptides was confirmed to be greater than 95% by high-performance liquid chromatography (HPLC), and the molecular weight of each peptide was verified by mass spectrometry (MS).

#### Confocal Microscopy

HeLa cells were maintained in DMEM supplemented with 10% fetal bovine serum (FBS) at 37°C in a humidified atmosphere containing 5% CO_2_. For confocal microscopy, cells were seeded at 5× 10^4^ cells/well in 24-well plates containing 13-mm round coverslips. Peptides labeled with 6-FAM were added to the culture medium at a final concentration of 10 *µ*M in a total volume of 500 *µ*L per well, and the cells were incubated for 1 hour at 37°C. After incubation, cells were fixed with 4% paraformaldehyde in PBS for 15 min at room temperature, permeabilized with 0.2% Triton X-100 in PBS for 15 min at room temperature, and blocked with 2% BSA in PBS for 30 min at room temperature. Cells were then stained for 1 hour at room temperature with either a rabbit anti-CXCR4 primary antibody (ab124824, Abcam) or a rabbit anti-NRP1 primary antibody (ab81321, Abcam), each diluted 1:1,000, followed by incubation for 1 hour at room temperature with a goat anti-rabbit IgG (H+L), cross-adsorbed, Alexa Fluor 594-conjugated secondary antibody (A-11012, Invitrogen) diluted 1:1,000. Cells were washed three times with PBS after fixation, permeabilization, blocking, and each antibody incubation step. Nuclei were counterstained using ProLong Gold Antifade Mountant with DAPI (P10144, Invitrogen).

Images were acquired using an EVIDENT FV4000 confocal laser scanning microscope with a UPLXAPO60XO 60× /1.42 NA objective lens. DAPI, 6-FAM, and Alexa Fluor 594 signals were acquired using the 405-, 488-, and 561-nm laser lines, respectively.

#### Quantification of Peptide Enrichment

Quantitative image analysis was performed in Fiji using a custom macro script. The main image-processing parameters are summarized in Table S2. The mean peptide fluorescence intensity was measured within the receptor-positive region (*I*_Receptor_) and within the peptide-positive region (*I*_Peptide_). The enrichment score was defined as

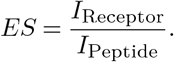

The final enrichment score for each image was obtained by averaging the four quadrant-level values. Full implementation details are provided in the Fiji macro available in the GitHub repository listed in the Data availability section. Representative raw images together with the corresponding peptide-positive and receptor-positive masks for Penetratin and S-02 are shown in Figs. S5 and S6, respectively.

## Supplementary Figures

**Figure S1.**
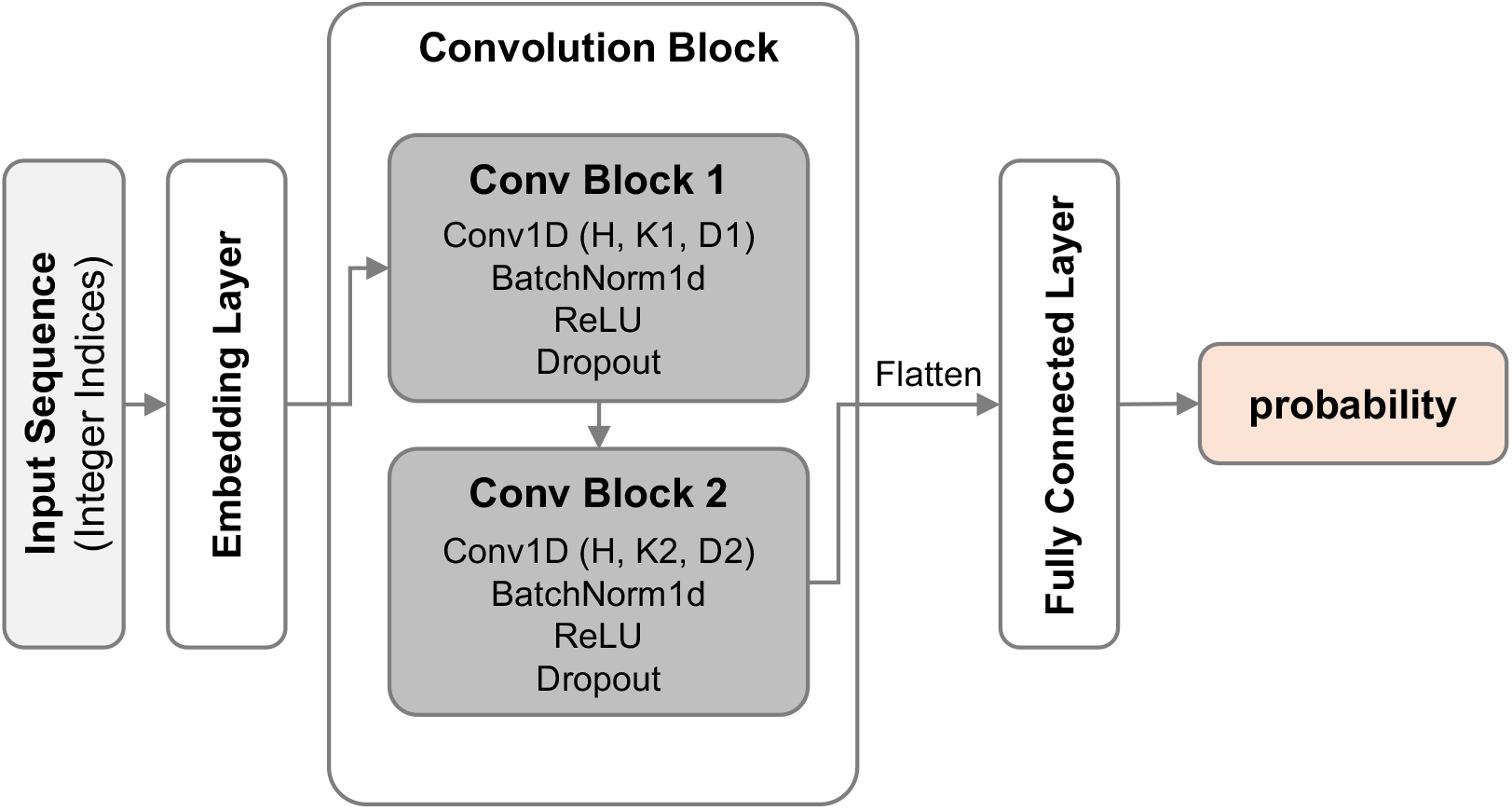
Architecture of the CPP binary classifier used in this study. Schematic illustration of the 1D-CNN model used to evaluate whether the generated sequences possess CPP-like characteristics. Input sequences were integer-encoded using a 20-amino-acid vocabulary with an additional PAD token, right-padded to the maximum sequence length observed in the development dataset, and passed through an embedding layer. The model consists of two 1D convolutional layers followed by batch normalization and dropout, a flattening layer, and a fully connected output layer followed by a sigmoid activation. Here, H denotes the number of convolutional output channels, and K1 and K2 denote the kernel sizes of the first and second convolutional layers, respectively. In this classifier, the embedding dimension and hidden dimension were both set to 16, the kernel sizes were 3 and 5, the dilation factor was fixed to 1 for both layers, and the dropout rate was 0.2.

**Figure S2.**
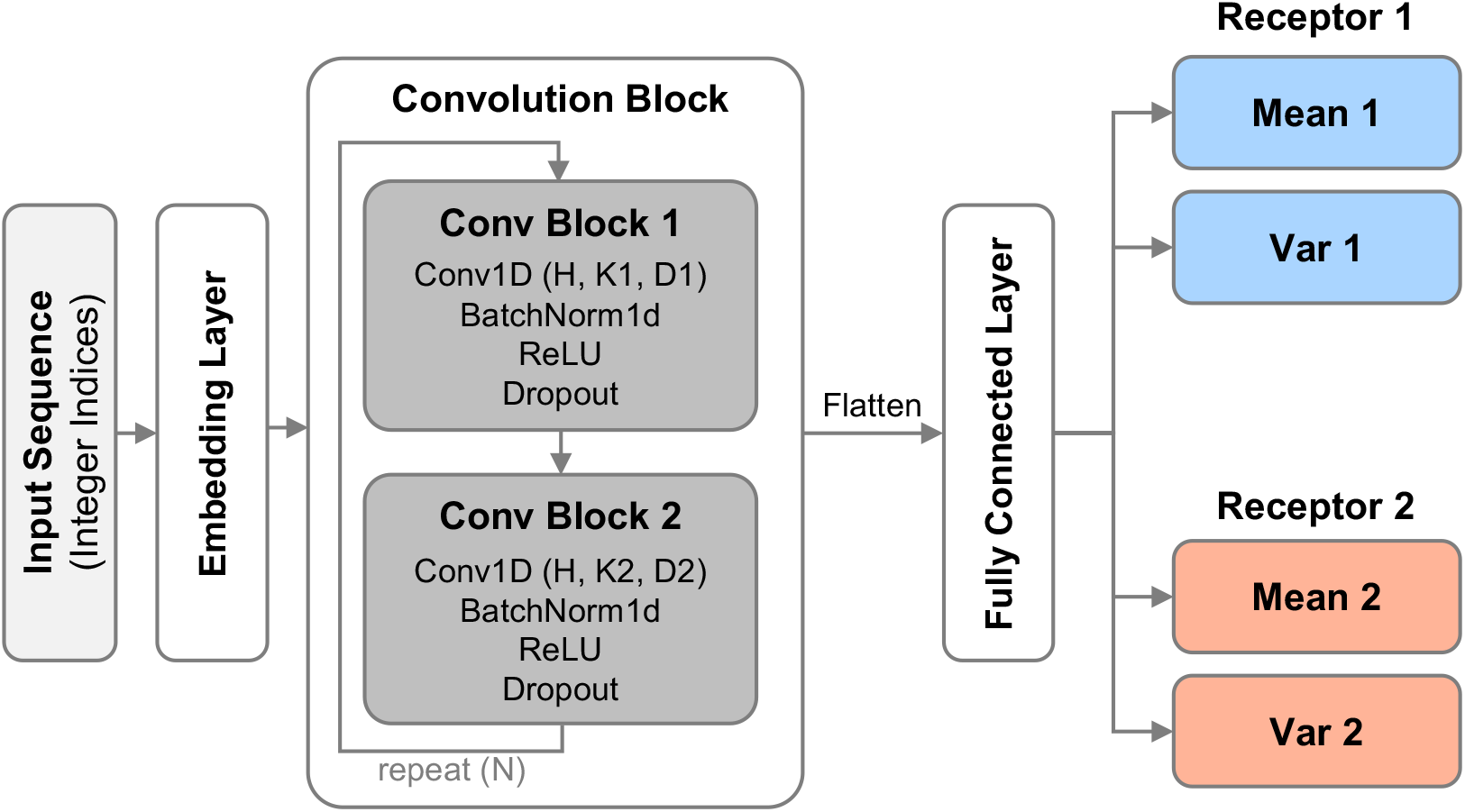
Architecture of the deep ensemble surrogate model. Schematic illustration of the convolutional regression model used to predict receptor-wise binding scores. For each receptor, the model outputs the predicted mean (*µ*) and variance (*σ*^2^) of the binding score. Convolutional layers extract local sequence patterns, which are then flattened and passed to regression heads that generate these outputs. Here, H denotes the hidden dimension (number of convolutional output channels), K1 and K2 denote the kernel sizes of the first and second convolutional layers, and D1 and D2 denote the corresponding dilation factors. The figure shows the common model structure shared across the ensemble, while architecture-specific hyperparameter settings are summarized in Table S1.

**Figure S3.**
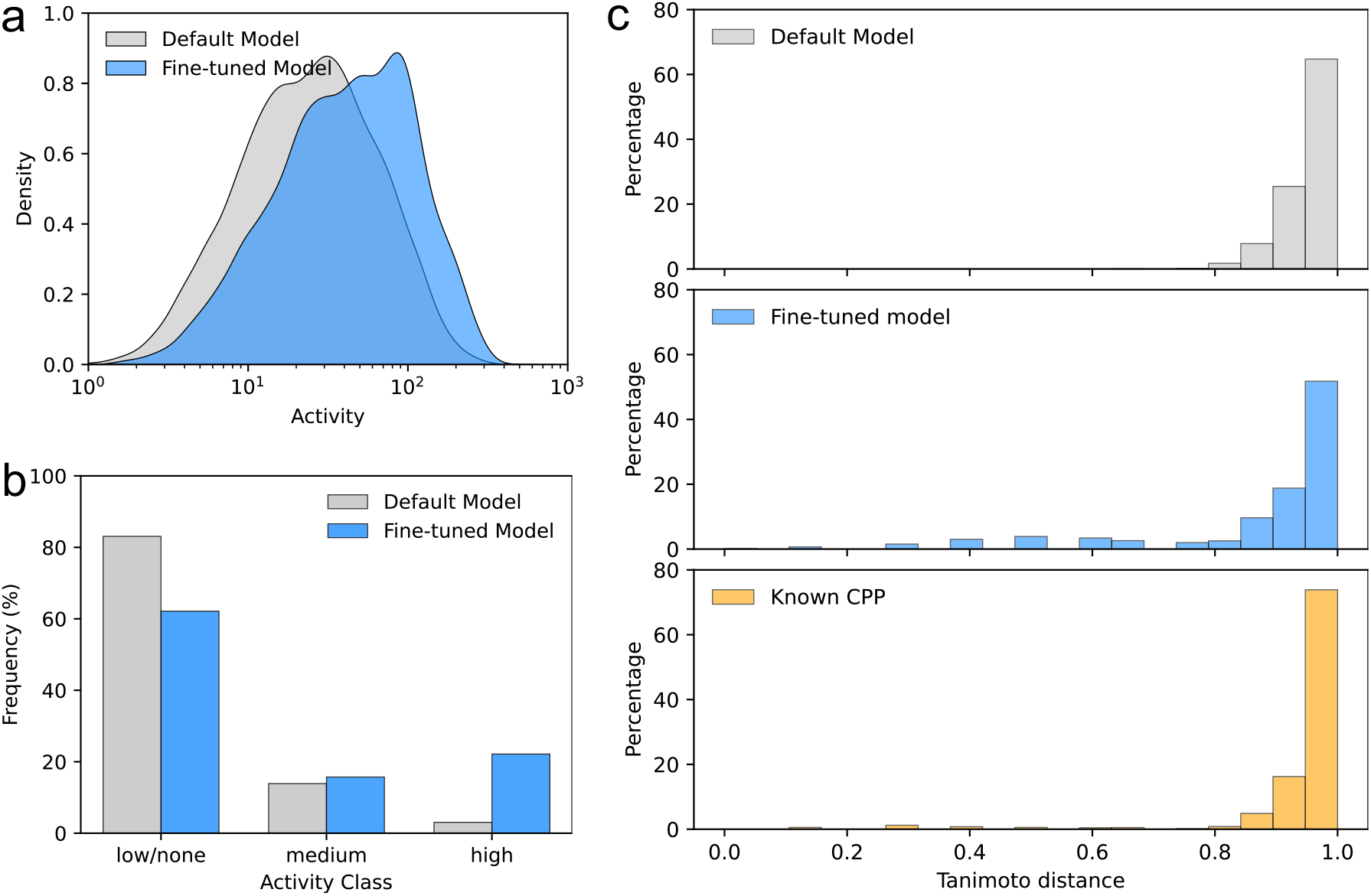
Evaluation of generated sequence characteristics. (a) Distribution of continuous CPP activity scores predicted by c3pred. (b) Activity class predictions by c3pred. (c) Histograms showing the distribution of pairwise Tanimoto distances for sequences, calculated on binary sequence fingerprints constructed by concatenating position-wise one-hot encodings of amino-acid identities. Top: sequences from the default EvoDiff model. Middle: sequences from the fine-tuned model. Bottom: known CPP sequences (training data).

**Figure S4.**
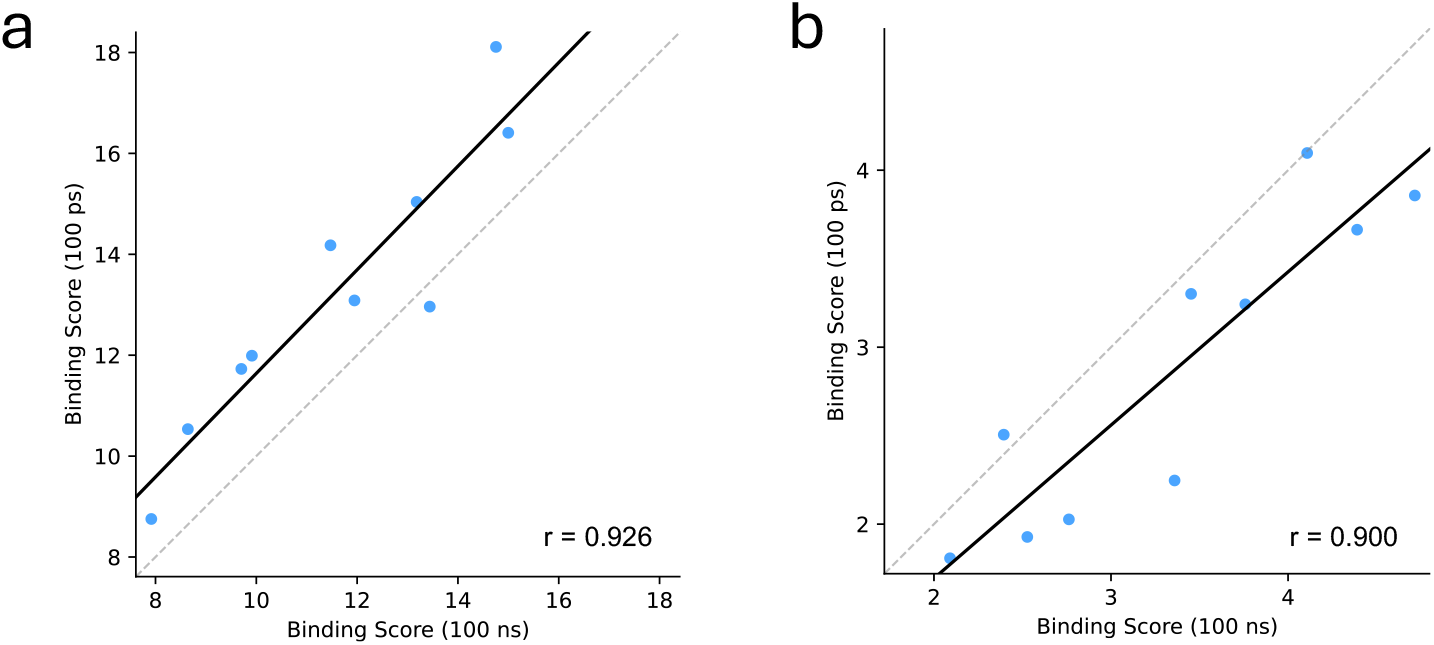
Comparison of binding scores between the 100 ps screening condition and the 100 ns reference condition. Binding scores obtained from the 100 ps screening condition and the 100 ns reference condition for the 10 peptides selected from the candidates evaluated during the optimization loop (S-01 to S-10). Each point represents one peptide, using scores averaged across simulations initiated from different initial structures for each peptide-receptor pair. (a) CXCR4. The 100 ns simulations were performed in a membrane-embedded system, whereas the 100 ps simulations were performed in water. Pearson’s correlation was *r* = 0.926 (*p* = 1.22 × 10^−4^), and Spearman’s correlation was *ρ* = 0.903. (b) NRP1. Both the 100 ps and 100 ns simulations were performed in water. Pearson’s correlation was *r* = 0.900 (*p* = 3.88 × 10^−4^), and Spearman’s correlation was *ρ* = 0.879.

**Figure S5.**
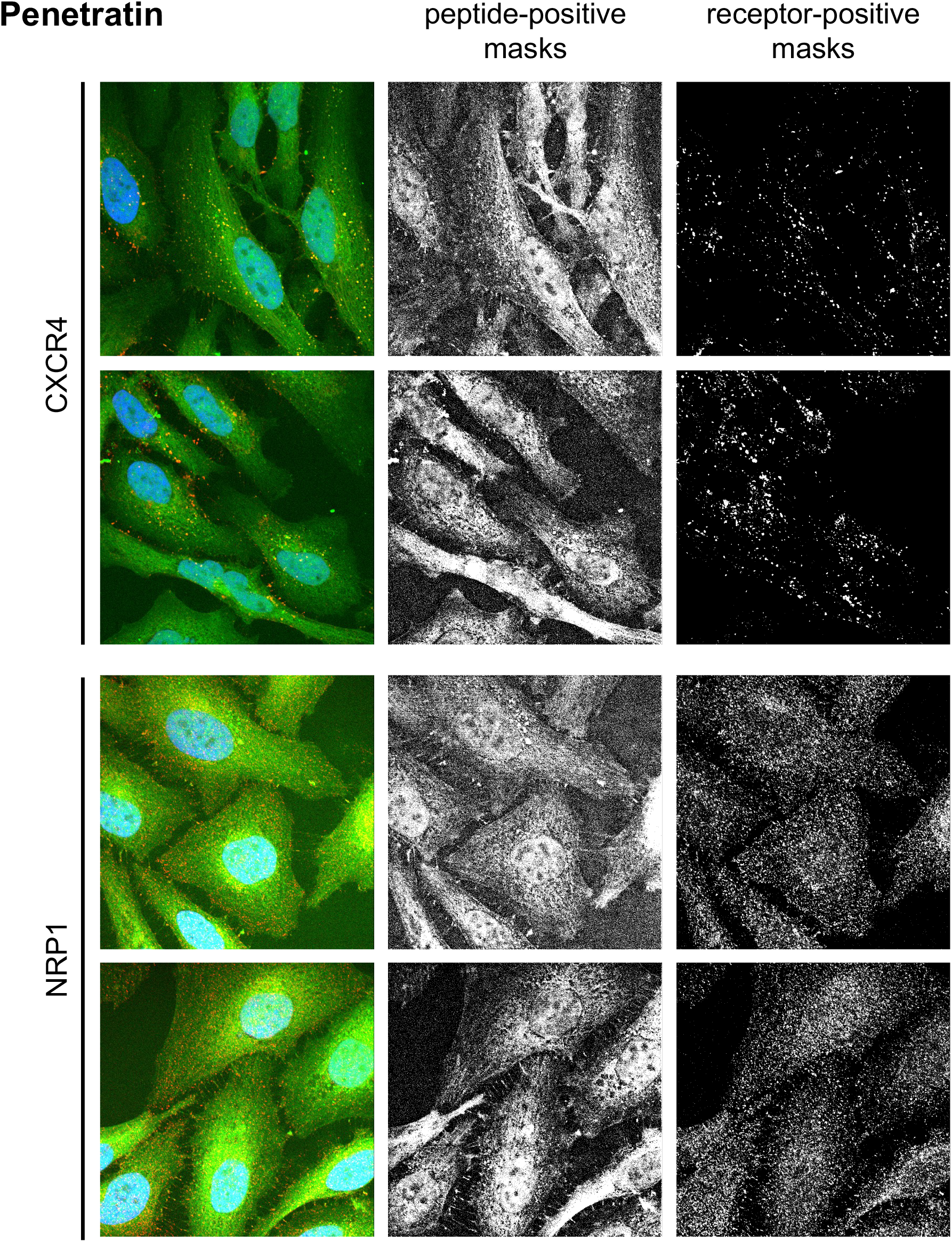
Representative confocal images and segmentation masks for Penetratin. HeLa cells were treated with 6-FAM-labeled Penetratin and immunostained for CXCR4 or NRP1. Top two rows: CXCR4 condition. Bottom two rows: NRP1 condition. Left: raw confocal images (Green: Penetratin; Red: receptor; Blue: DAPI). Center: peptide-positive masks. Right: receptor-positive masks.

**Figure S6.**
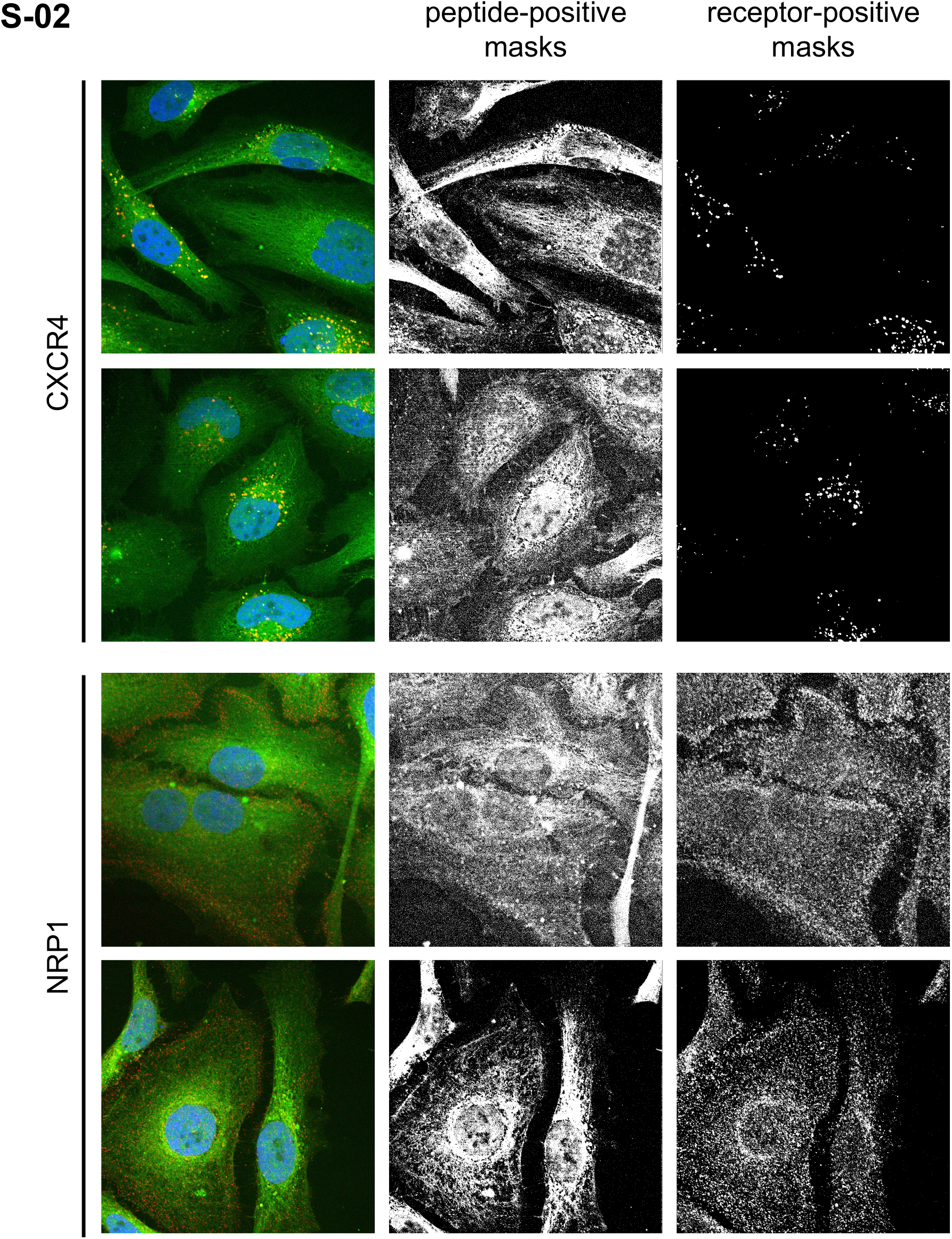
Representative confocal images and segmentation masks for peptide S-02. HeLa cells were treated with 6-FAM-labeled S-02 and immunostained for CXCR4 or NRP1. Top two rows: CXCR4 condition. Bottom two rows: NRP1 condition. Left: raw confocal images (Green: S-02; Red: receptor; Blue: DAPI). Center: peptide-positive masks. Right: receptor-positive masks.

**Figure S7.**
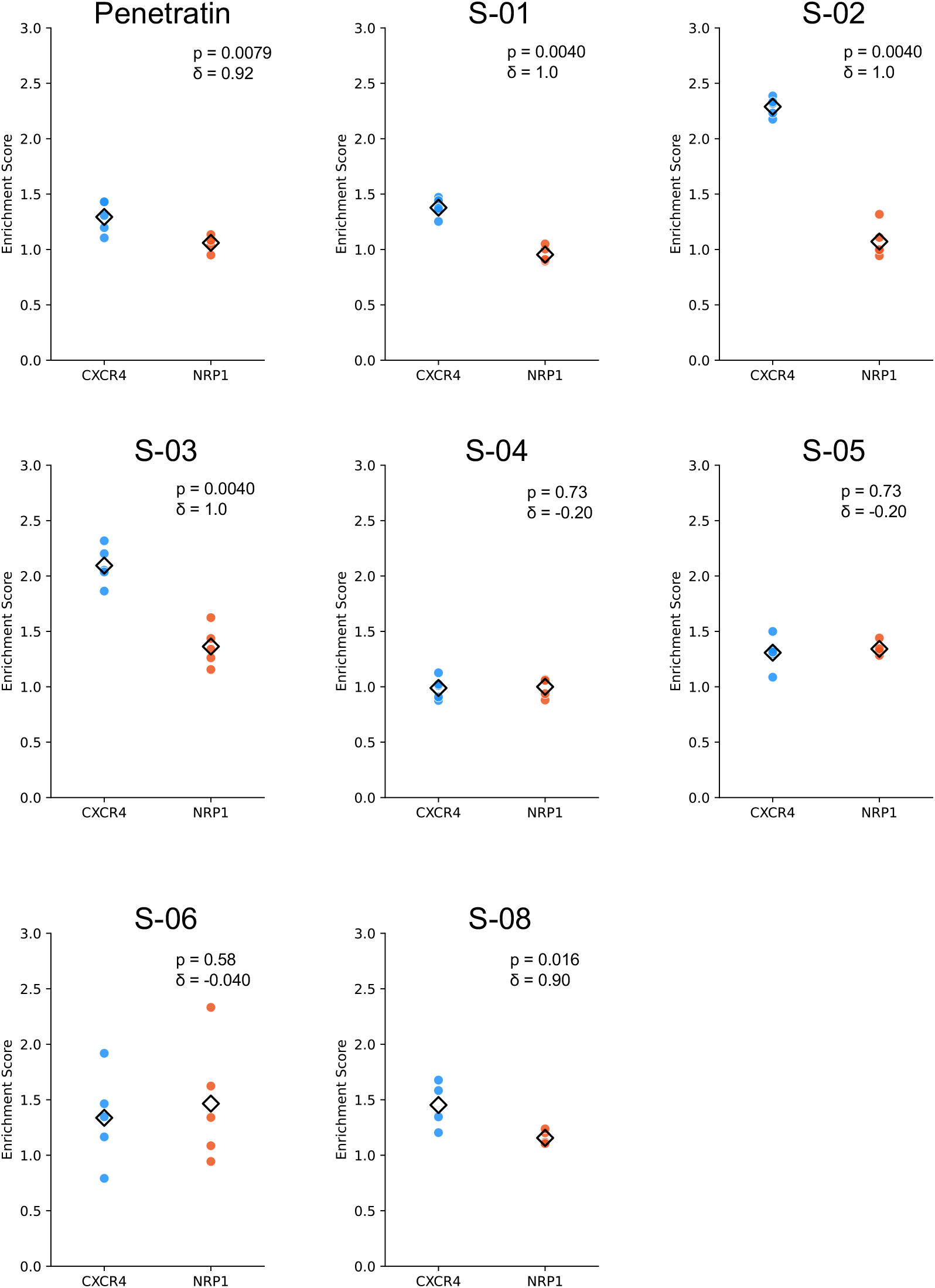
Receptor-associated enrichment scores for selected peptides. Quantitative comparison of enrichment scores for Penetratin and selected peptides (S-01 to S-06, S-08). TAT, R8, S-07, S-09, and S-10 were excluded because fluorescence intensity was insufficient for reliable quantification. Each dot represents a single field of view, and diamonds indicate the mean. Panel annotations show one-sided Mann-Whitney *U* test *p* values for the comparison CXCR4 > NRP1 and Cliff’s *δ*.

**Table S1.**
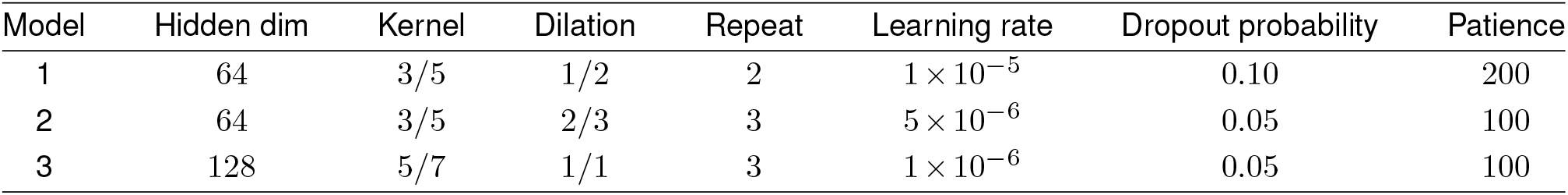
Architecture and training hyperparameters of the deep ensemble models.

**Table S2.**
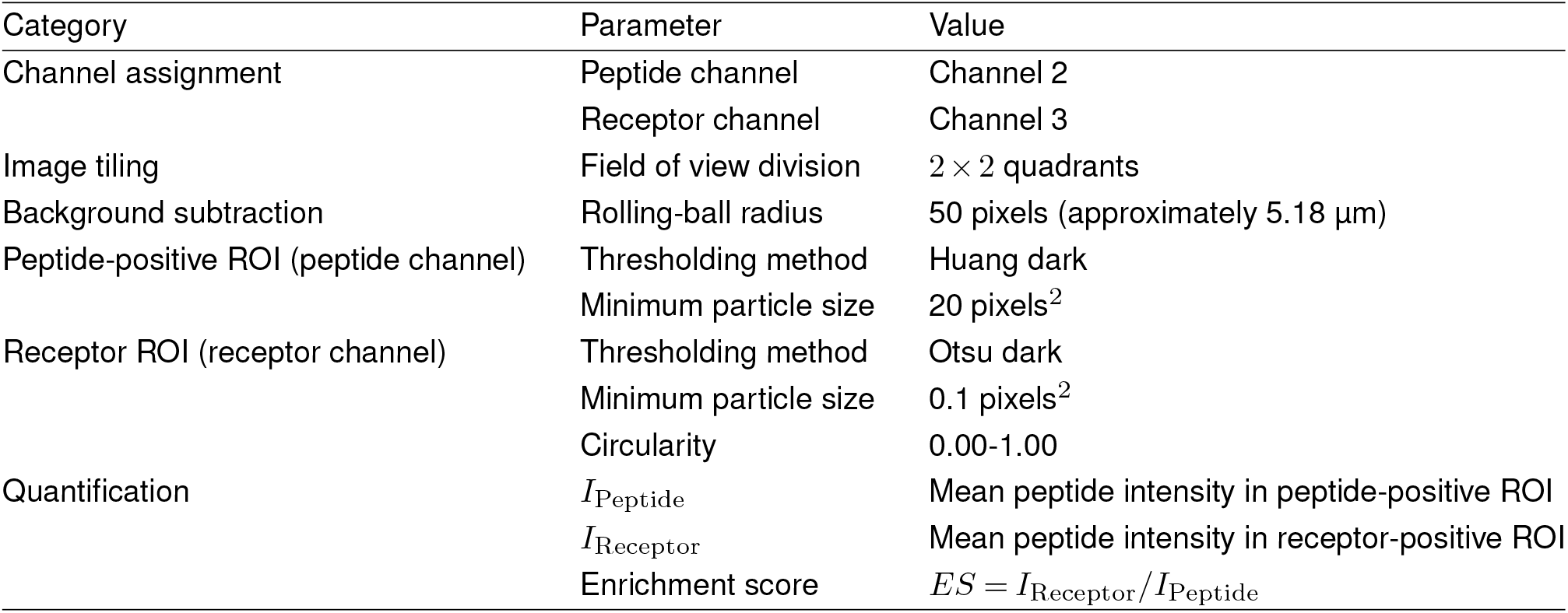
Image-processing parameters used for enrichment-score quantification in Fiji.

| Category | Parameter | Value |
| --- | --- | --- |
| Channel assignment | Peptide channel | Channel 2 |
|  | Receptor channel | Channel 3 |
| Image tiling | Field of view division | $2 \times 2$ quadrants |
| Background subtraction | Rolling-ball radius | 50 pixels (approximately 5.18 $\mu\text{m}$) |
| Peptide-positive ROI (peptide channel) | Thresholding method | Huang dark |
|  | Minimum particle size | 20 pixels<sup>2</sup> |
| Receptor ROI (receptor channel) | Thresholding method | Otsu dark |
|  | Minimum particle size | 0.1 pixels<sup>2</sup> |
|  | Circularity | 0.00-1.00 |
| Quantification | $I_{\text{Peptide}}$ | Mean peptide intensity in peptide-positive ROI |
| | $I_{\text{Receptor}}$ | Mean peptide intensity in receptor-positive ROI |
| | Enrichment score | $ES = I_{\text{Receptor}}/I_{\text{Peptide}}$ |

## Notes

### Competing Interest Statement

The authors have declared no competing interest.

